# Ketamine decreases HPA axis reactivity to a novel stressor in male but not female mice

**DOI:** 10.1101/2021.06.29.450387

**Authors:** Colin J. Johnston, Paul J. Fitzgerald, Jena S. Gewarges, Brendon O. Watson, Joanna L. Spencer-Segal

**Affiliations:** Michigan Neuroscience Institute, University of Michigan, Ann Arbor, MI; Department of Psychiatry, University of Michigan, Ann Arbor, MI; Neuroscience Graduate Program, University of Michigan; Department of Internal Medicine, Division of Metabolism, Endocrinology, and Diabetes, University of Michigan, Ann Arbor, MI

**Author notes:** To whom correspondence should be sent (205 Zina Pitcher Place, Ann Arbor MI, 48109). Declarations of interest: none.

**Keywords:** ketamine, metyrapone, fecal corticosterone, forced swim test, chronic stress, depression

## Abstract

Ketamine is an antidepressant drug that interacts with the hypothalamic-pituitary-adrenal (HPA) axis, but whether this interaction is important for its behavioral effect is unknown. The goal of this experiment was to determine whether the behavioral response to ketamine depends on intact HPA axis function. Male and female C57BL/6J mice underwent chronic unpredictable stress prior to ketamine (30 mg/kg, i.p.) or vehicle treatment, with or without the glucocorticoid synthesis inhibitor metyrapone (20 mg/kg, i.p.) to block adrenal corticosterone production. Mice were tested in the forced swim test (FST) and open field test one and two days after injection, respectively. Fecal corticosterone was measured at select time points. No significant drug effects on behavior were observed. Males consistently had higher fecal corticosterone levels and stress-induced increases than females. Ketamine lowered the fecal corticosterone response to the FST only in males. These data show that ketamine after chronic stress decreases the corticosterone response to a novel stressor (the FST) in males, but not females. Corticosterone levels in all mice correlated with immobility in the FST, suggesting that shared neural circuitry could mediate both endocrine and behavioral responses. This circuitry may be ketamine-responsive only in males.

## 1. Introduction

Ketamine has been extensively studied as a rapid-acting antidepressant in humans and rodents. In humans, ketamine is used for the treatment of refractory Major Depressive Disorder, a stress-related psychiatric disease. In rodents, ketamine may be more effective when the subjects had previously been chronically stressed (Fitzgerald et al., 2019). Because of the relationship of stress to depression and ketamine action, we hypothesized that the antidepressant effect of ketamine could occur via its effect on the neuroendocrine stress response via the hypothalamic-pituitary-adrenal (HPA) axis.

Ketamine has both acute and chronic effects on HPA axis activity. For example, ketamine infusion acutely increased corticosterone in rats (Radford et al., 2018) and cortisol in humans (Khalili-Mahani et al., 2015). On the other hand, ketamine reversed chronic stress-induced HPA axis hyperactivity in rats (Garcia et al., 2009), and in one human case report (Ostroff & Kothari, 2015). These studies indicate that ketamine affects acute and chronic HPA axis activity differently and that these mechanisms may be conserved between rats and humans.

Based on these previous studies, we hypothesized that ketamine would increase corticosterone acutely, and that ketamine would normalize subsequent HPA axis hyperactivity in chronically stressed male and female mice. Ketamine’s antidepressant activity is thought to be due, at least in part, to its ability to antagonize NMDA receptor signaling. (Fuchikami et al., 2015; Klein et al., 2020; Sleigh et al., 2014; Williams et al., 2018; Yang et al., 2018; Zanos et al., 2016). Since corticosterone also influences NMDA signaling (Mikasova et al., 2017), we hypothesized that ketamine’s behavioral efficacy would depend on corticosterone availability at the time of injection. To study this, we blocked adrenal steroidogenesis using metyrapone prior to ketamine injection in male and female mice with a history of chronic unpredictable stress and studied the downstream effect on behavior and fecal corticosterone.

## 2. Methods

### 2.1 Subjects

40 male and 40 female C57BL/6J 10-12 week old mice (The Jackson Laboratory, Bar Harbor ME) were single-housed as previously described; the sample size was justified through the observed effects of the previous experiment (Fitzgerald et al., 2019). Experimental protocols were approved by the University of Michigan Institutional Animal Care and Use Committee and conducted in accordance with the NIH Guide for the Care and Use of Laboratory Animals.

These experiments were carried out in two cohorts, segregated by sex. A locotest was performed on arrival. Each mouse was allowed to explore a 36 cm square arena for 5 minutes at low light (30 lux). Mice were placed into groups with equivalent average baseline locomotion. Two weeks later, the stress group underwent chronic unpredictable stress (continued single housing, minimal handling and cage enrichment, and a daily hassle as previously described (Fitzgerald et al., 2019)). Unstressed mice were handled for 30 seconds daily for the first 5 days, and had cage enrichment (Fitzgerald et al., 2019). On the day after the final stressor, mice underwent two intraperitoneal injections 45 minutes apart. The first injection contained either vehicle (0.9% saline) or the 11-beta hydroxylase inhibitor metyrapone (20 mg/kg) (Fisher Scientific) in saline solution. The second contained either vehicle or 30 mg/kg (*R,S*)-ketamine (Par Pharmaceutical) in saline. The experiment included the following groups (n = 8 mice/sex/group): no stress/vehicle/vehicle, stress/vehicle/vehicle, stress/metyrapone/vehicle, stress/vehicle/ketamine, and stress/metyrapone/ketamine. The day after injection, mice underwent forced swim testing (FST), and on the following day the open field test (OFT) (Figure 1a).

**Figure 1.**
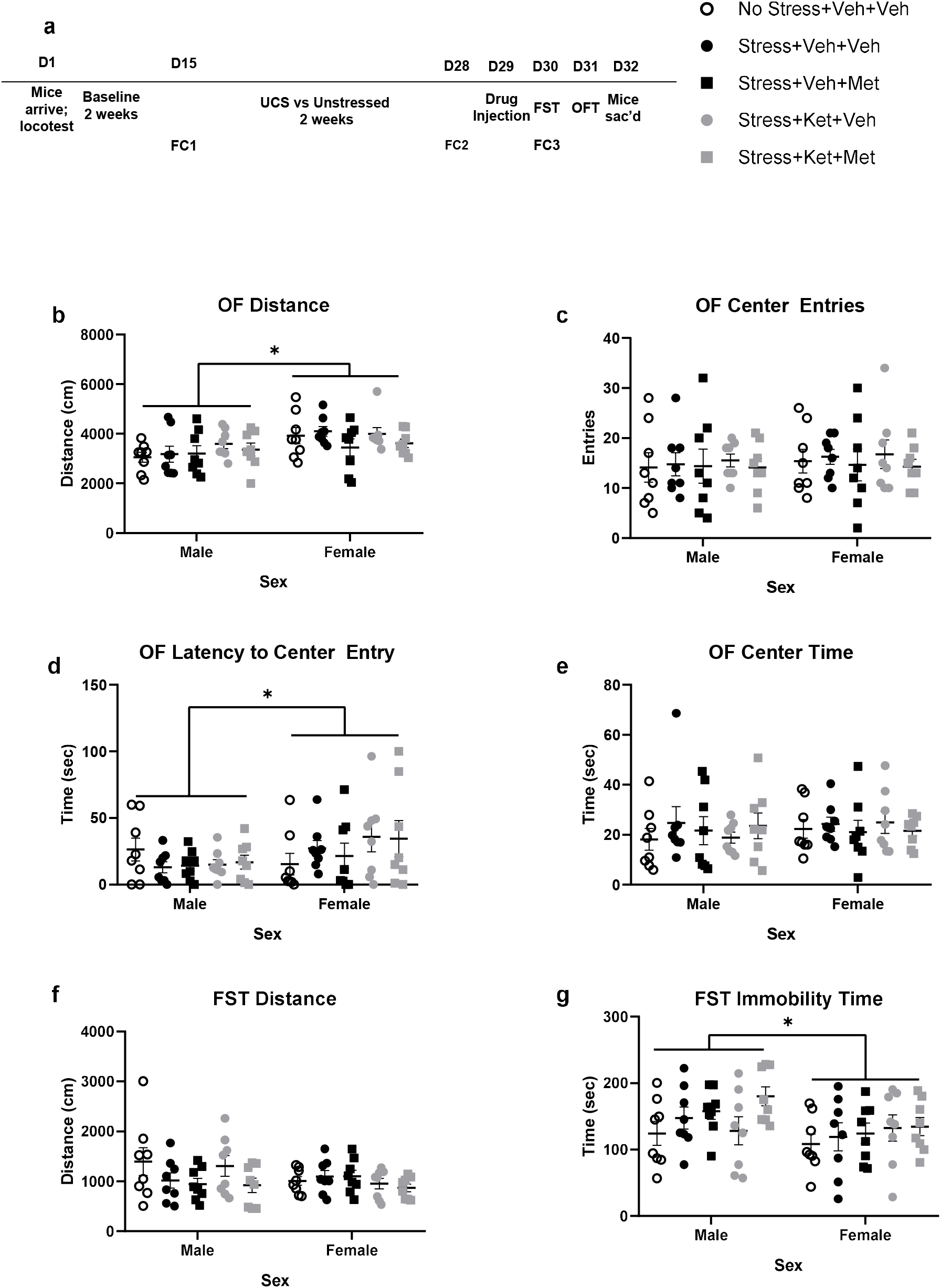
Sex differences in behavior in the FST and OFT. Behavior was not significantly modulated by ketamine or stress, but male mice given ketamine trended toward an antidepressant-like effect in the FST that was counteracted by metyrapone. Experimental timeline (a), behavior measurements for the OF (b-e) and FST (f-g) separated by sex and treatment groups. Data are shown as mean ±SEM. * *p* < 0.05.

For all behavioral tests, mice were acclimated to the testing room for at least 30 minutes prior to the start. Testing was performed during the light phase. Videos were analyzed with Ethovision version 11.5.

### 2.2 Forced Swim Test

The forced swim apparatus was 4 cylindrical tanks 20 cm in diameter separated by opaque barriers. The tanks were halfway filled with water at 23-25°C. Mice were placed one per tank for 6 minutes at 200 lux. The mouse was immediately dried off afterward and returned to its homecage. After 5 tests, the water was changed, and the tanks were wiped down with 50% ethanol. For behavior analysis, mobility was defined as at least 2 cm/s, and immobility was defined as at most 1.75 cm/s.

### 2.3 Open Field Test

The open field apparatus was a 72 square arena with walls 36 cm tall and its center defined as the center square with sides of 36 cm. Each mouse was allowed to explore the open field for 5 minutes, at 200 lux. The open field apparatus was cleaned with 50% ethanol between animals.

### 2.4 Fecal Collection and ELISA Preparation

Corticosterone was measured from feces at 3 time points in order to avoid the stress of repeated blood collection. Each collection (FC) was performed at 3pm. FC1 took place after the two-week baseline period, before the stress protocol began. FC2 took place after stress, the day prior to injections. Cages were changed at 9am the day of these two collections. FC3 took place after the FST to measure the effect of this acute stressor. Mice were placed into a clean cage immediately after swimming, and all feces were collected for the next 6 hours. Fecal pellets were frozen at -20°C until analysis. Defrosted samples were desiccated for 3 days, pulverized, weighed, diluted with 5μL of 100% ethanol per mg of sample and shaken overnight. Samples were centrifuged at 4°C for 15 minutes at 5000 rpm, and the supernatant saved. 100 μL of each sample was desiccated. The day of each ELISA, 200 μL of each sample was reconstituted as described in the ELISA kit. Arbor Assays DetectX Corticosterone ELISA kit (K014-H) was used to measure corticosterone.

### 2.5 Statistical Analysis

For all behavioral variables, an unpaired two-tailed t-test was used to compare the no stress group with the stressed vehicle group. For fecal corticosterone, a paired two-tailed t-test was used to compare FC1 to FC2, to see if the chronic variable stress paradigm alone had any effect on fecal corticosterone. FC3 was then normalized to FC2 per subject (nFC3). For the remaining analyses, three-way between-subjects ANOVAs (ketamine by metyrapone by sex) were conducted using all stressed animals, as well as two-way ANOVA within each sex, with post-hoc tests as appropriate with Bonferroni corrections. Statistical significance was defined as *p* < 0.05. Pearson correlation was used to assess correlation of nFC3 with immobility in the FST.

## 3. Results

### 3.1 Behavior

We observed sex differences in behavior (Figures 1b-g). In the OFT, females travelled farther than males (*F*(1,56f) = 6.025, *p*=.017) and entered the center later (*F*(1,56) = 7.122, *p*=.010). In the FST, males were immobile for longer than females (*F*(1,56) = 4.572, *p*=.038). There were no significant differences in behavior based on stress or drug treatment.

For males and females separately, stress again had no significant effect on behavior for either sex. For the males, there were no effects of ketamine or metyrapone in the stressed groups. For the females, a main effect of metyrapone on open field distance was observed, such that females given metyrapone travelled less than those given vehicle (*F*(1,28) = 4.337, *p*=.047).

While no significant drug effects were observed on behavior, males given ketamine trended toward reducing depressive-like behaviors in the FST, while the presence of metyrapone seemed to abolish this effect (Figure 1g). No trends were observed in females.

### 3.2 Corticosterone Measurements

We conducted noninvasive measures of corticosterone from feces at multiple time points to understand the effect of our stress paradigm and ketamine treatment on basal and swim-stimulated corticosterone (Figures 2a-c). Fecal corticosterone measurements were used to avoid the additional stress of blood collection and to capture the total activation and termination of the stress response after specific manipulations (see Methods). We collected fecal corticosterone at three timepoints (FC1-3).

**Figure 2.**
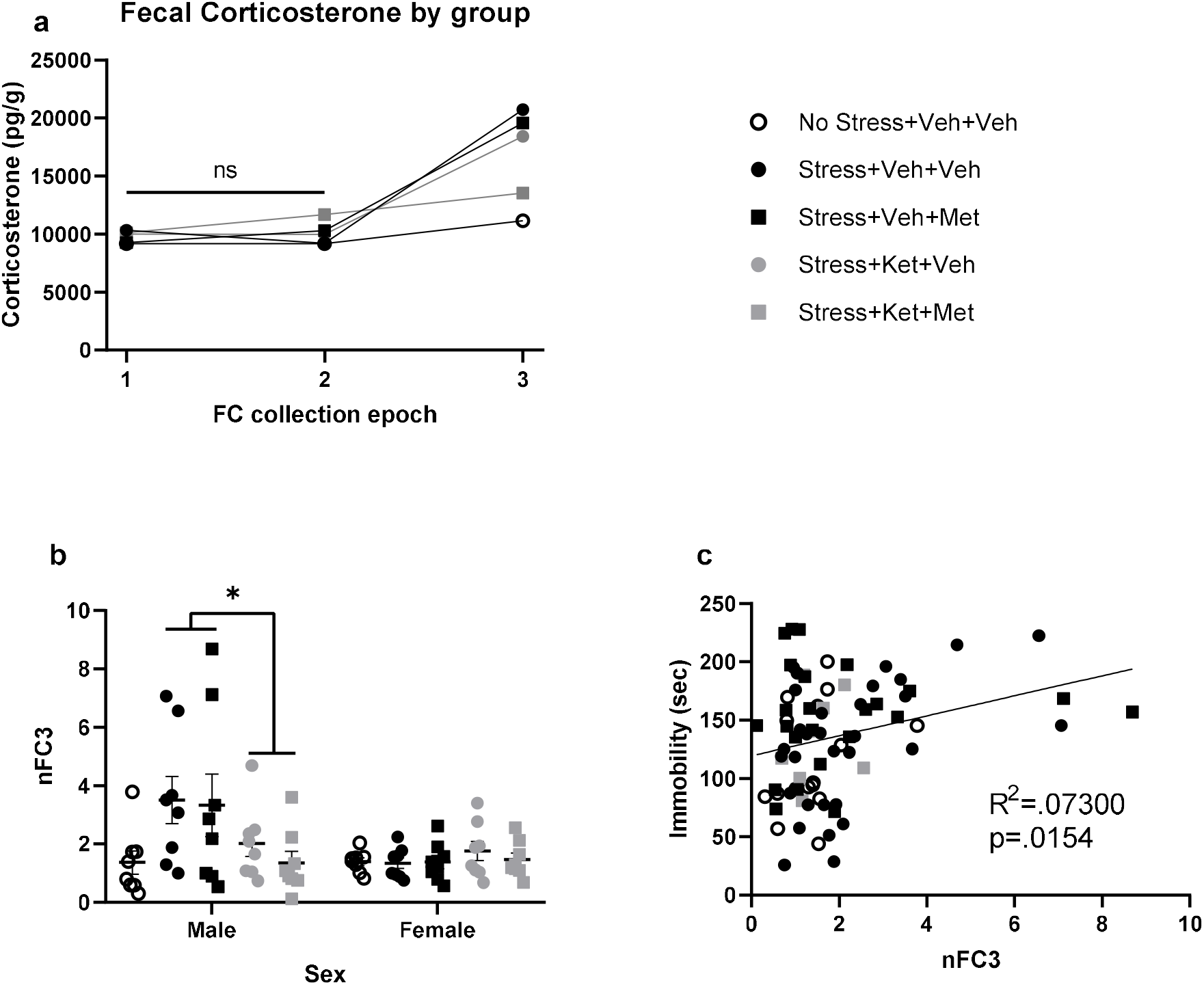
Prior ketamine normalizes post-swim corticosterone in male but not female mice. a) Raw fecal corticosterone (FC) values by group; b) Normalized FC3 measurements for males and females; and c) Correlations between nFC3 and immobility in the FST in both sexes combined. Data are shown as mean ± SEM. * *p* < 0.05.

FC2 (post-chronic stress) was not significantly different from FC1 (prior to stress) for either the non-stressed (*t*(15) = 0.013, *p*=.99) or the stressed groups (*t*(63) = 0.394, *p*=.70) (Figure 2a), indicating no effect of the stress paradigm on baseline corticosterone. We therefore normalized each animal’s FC3 and to FC2 division to reflect a change relative to FC2, thereby creating normalized value: nFC3, which represents the fecal corticosterone after forced swimming relative to the baseline collection.

Males had higher nFC3 than females *F*(1,56) = 7.607, *p*=.008). This was also true for the non-normalized FC3 (*F*(1,55) = 14.804, *p* < 0.001) values (Figure 2b).

For nFC3, there was an interaction between sex and ketamine (*F*(1,56) = 6.598, *p*=.013), with post-hoc tests showing that ketamine decreased nFC3 in males (*t*(30) = 2.432, *p*=.042), but not in females (*t*(30) = 1.039, *p*=.61) (Figure 2c). Furthermore, stress on its own caused significantly higher nFC3 in males (*t*(14)=2.389, *p*=.03). Similarly, two-way ANOVA of nFC3 for stressed males and females separately, showed a main effect of ketamine on nFC3 only in males (*F*(1,28) = 5.607, *p*=.025). Thus, in males only, prior chronic stress increases the corticosterone response to forced swimming, and ketamine normalizes this.

In a Pearson correlation using all the data, higher immobility in the FST correlated with higher nFC3 (*r*(78) = 0.07300, *p*=.015) (Figure 2d).

## 4. Discussion

In this study we tested the effects of ketamine HPA axis activity in male and female mice. We also tested whether the behavioral efficacy of ketamine depended on intact adrenal corticosterone synthesis at the time of injection. Our interpretation of the behavioral interaction between ketamine and metyrapone is limited by the lack of significant effect of stress or ketamine on behavior. While we did not see clear behavioral effects of ketamine, we did find that stressed male mice showed HPA axis hyperactivity to the FST, a novel stressor, and that ketamine normalized this response.

The lack of a significant behavioral effect of stress or ketamine shown here contrasts with other published studies (Fitzgerald et al., 2019; Podkowa et al., 2016; Wang et al., 2011). There was, however, a trend of behavior in males as hypothesized, such that stressed males tended to exhibit higher immobility in the FST, with ketamine and metyrapone tending to decrease or increase immobility, respectively. This did not occur in females. One study that reported decreased immobility in the ketamine-treated mice of both sexes after stress (Franceschelli et al., 2015) used lower doses of ketamine than the doses used in this study (3-10 vs 30 mg/kg), and females were more sensitive to the lower doses than males. Thus, the behavioral effects of ketamine after stress may be dose-dependent, especially in females. It is also possible that the stress paradigm was less efficacious in females.

In keeping with the possibility of greater stress and ketamine sensitivity in males, we observed that stressed males had higher corticosterone after the FST than nonstressed males, while females did not, and ketamine normalized this corticosterone response in males. This is consistent with a previous study in which ketamine reduced circulating corticosterone after chronic stress in male rats (Garcia et al., 2009). Reasons for this sex difference could include lack of sensitivity of females to our stress paradigm and/or to ketamine at this dose. Our observation of higher fecal corticosterone levels and higher fecal corticosterone increases in males contrasts with the higher circulating corticosterone and higher circulating corticosterone increases in females, and may be explained by the higher level of corticosterone binding globulins and lower free corticosterone seen in female rodents (Minni et al., 2014).

The relevance of post-FST corticosterone is demonstrated by its positive correlation with FST immobility on a per-animal basis, when the two sexes were combined. The correlation suggests that shared neural circuitry could mediate endocrine and immobility responses to the FST. While ketamine did not significantly change the behavior of male mice in this study, the correlation between corticosterone and immobility and our finding of decreased corticosterone in ketamine-treated male mice is consistent with the often reported finding that ketamine decreases FST immobility (Podkowa et al., 2016).

In conclusion, we showed that ketamine normalizes HPA axis hyperreactivity to a novel stressor in stressed male mice. In male mice, the corticosterone response to a novel stressor may be a biomarker of ketamine’s effect. Future studies should determine whether the sex differences in ketamine’s actions are qualitative, representing different physiologic mechanisms or neural circuitry, or quantitative, representing differential sensitivity or dose-response.

